# Design of a transcriptional biosensor for the portable, on-demand detection of cyanuric acid

**DOI:** 10.1101/736355

**Authors:** Xiangyang Liu, Adam D. Silverman, Khalid K. Alam, Erik Iverson, Julius B. Lucks, Michael C. Jewett, Srivatsan Raman

## Abstract

Rapid molecular biosensing is an emerging application area for synthetic biology. Here, we engineer a portable biosensor for cyanuric acid (CYA), an analyte of interest for human and environmental health, using a LysR-type transcription regulator (LTTR) from *Pseudomonas* within the context of *Escherichia coli* gene expression machinery. To overcome cross-host portability challenges of LTTRs, we rationally engineered hybrid *Pseudomonas-E. coli* promoters by integrating DNA elements required for transcriptional activity and ligand-dependent regulation from both hosts, which enabled *E. coli* to function as whole-cell biosensor for CYA. To alleviate challenges of whole-cell biosensing, we adapted these promoter designs to function within a freeze-dried *E. coli* cell-free system to sense CYA. This portable, on-demand system robustly detects CYA within an hour from laboratory and real-world samples and works with both fluorescent and colorimetric reporters. This work elucidates general principles to facilitate the engineering of a wider array of LTTR-based environmental sensors.

## INTRODUCTION

Rapid molecular biosensing is an emerging application area for synthetic biology, particularly for the on-demand detection of viruses^1–5^ microbes^6,7^, metals^8,9^, and pollutants^10,11^ in biological or environmental samples. Traditional methods for detecting these analytes of interest require collection of large-volume samples on-site and transportation to centralized laboratories equipped with sophisticated analytical equipment. These approaches are costly, time-consuming, and require extensive infrastructure and technical expertise that may not be readily available in resource-limited or time-sensitive settings. In contrast, biosensors engineered to be sensitive and specific for particular analytes could be deployed directly for molecular detection at the site of sampling. For this reason, biosensors offer great appeal for rapid and scalable molecular detection.^12^

Biosensors are typically designed by identifying natural biorecognition elements and then engineering them into a genetic circuit to drive the expression of a reporter gene in response to a desired target molecule.^12,13^ Synthetic biosensors can be deployed as whole cells, in which the circuit is embedded in live bacteria, or cell-free biosensors, in which the circuit functions wholly within a cell extract or *in vitro* reaction mixture. Although theoretically simple, the engineering of either whole-cell or cell-free biosensors is unpredictable and often requires significant optimization. Biorecognition elements are frequently allosteric transcription factors (aTFs), which interact with a cognate promoter through ligand-dependent conformational changes to regulate gene expression.^14^ However, aTFs and the promoters they regulate are often layered in complex networks of genetic regulation. Since these interactions cannot be easily disentangled, native promoters can be difficult to engineer for properties like fold-activation and sensitivity. Native promoters also exhibit context dependence, which complicates the porting of aTFs from non-model hosts (e.g., *Pseudomonas* or *Acinetobacter*) into common laboratory strains of *E. coli*.^15–17^ Thus, repurposing the large diversity of known aTFs and the molecules they sense into useful diagnostic tools requires a better understanding of their fundamental mechanism of action.

Even for aTFs that have successfully been integrated into functional whole-cell sensors (for example: lead,^18^ mercury,^19^ arsenic,^8^ phenanthrene,^20^ and trinitrotoluene^10^), practical challenges still remain for their use in on-demand field sample testing. These include the fact that whole-cell biosensors have to be encapsulated and maintained in conditions that promote bacterial survival in order to be functional. Furthermore, natural biological processes could result in plasmid loss, mutation, recombination, horizontal gene transfer, or other events that abrogate sensor function.^21^ Whole-cell, one-component sensors are also not viable for the detection of cytotoxic or membrane-impermeable analytes such as organophosphates or antibiotics. Finally, genetically-modified organisms are subject to biocontainment and regulatory scrutiny, which challenges their deployment in the field.

Cell-free biosensors show promise for solving some of the practical issues constraining whole cell biosensors by replacing cellular transcription and translation with cell-free gene expression (CFE) instead. A CFE reaction is a mixture of a processed bacterial lysate with amino acids, nucleotides, salts, cofactors, energy substrate, and DNA template(s) encoding the gene(s) to be transcribed and translated *in vitro*.^22^ CFE reactions offer a number of advantages relative to whole-cell sensors, particularly for use in environmental biosensing: they are not subject to growth and mutational constraints, they are less restricted by analyte toxicity and diffusion limitations, and they have reduced concerns for biocontainment. Additionally, recent work in the design of cell-free nucleic acid biosensors has demonstrated that cell-free reactions can retain functionality when stored lyophilized (freeze-dried) at ambient temperature for months.^1,2,23^

Here, we report the design of novel whole-cell and cell-free biosensors for detecting cyanuric acid (CYA) using AtzR, a CYA-responsive aTF. AtzR is a LysR-type transcription regulator (LTTR) found in *Pseudomonas* sp. strain ADP1.^24^ We chose AtzR as LTTRs are among the largest family of bacterial aTFs and any design principles we uncover could facilitate the engineering of new LTTR biosensors.^25,26^ Additionally, the mechanism by which LTTRs regulate transcription is complex and constitutes a challenging proof-of-principle for porting biosensing circuits into *E. coli*-based biosensors. In addition, CYA is an important environmental and human toxin. Widely used as a stabilizer for chlorine tablets, CYA must be kept at tightly controlled levels in swimming pools to prevent microbial growth. Most current methods for CYA detection utilize unreliable test strips or are based on an analysis of turbidity resulting from a non-specific CYA precipitation.

Previously, a CYA sensor was developed by utilizing native promoter of *atzDEF* from *Pseudomonas* sp. strain ADP1 in *E. coli* without optimizing for the non-native host.^27^ However, poor compatibility of native *Pseudomonas* promoter with *E. coli* transcriptional machinery gave low reporter signal and activation, and general design rules could not be inferred from the implementation. To overcome these shortcomings, we sought to use this system to develop a CYA biosensor in *E. coli* that would enable us to elucidate design rules for activating LTTRs in non-native hosts. We first rationally designed a library of hybrid promoters based on the native *Pseudomonas atzDEF* promoter and a constitutive *E. coli* σ70 promoter and identified a number of designs that were inducible by CYA. Systematic characterization of the whole-cell CYA biosensor with respect to timing and growth stage revealed that the sensor was functional, but practical challenges remained in deploying whole cell biosensors. To overcome these challenges, we therefore sought to develop a cell-free CYA biosensor by implementing the responsive promoters in *E. coli* extract CFE reactions. Characterization of the cell-free biosensor confirmed that it specifically responds to CYA when challenged with unfiltered environmental water samples, that it is compatible with both fluorescent and colorimetric reporters, and that it can also be lyophilized for long-term storage and distribution. This work represents the first implementation of an LTTR in a bacterial extract, expanding the range of aTFs that have been shown to be functional in freeze-dried, cell-free systems beyond the well-characterized transcriptional regulators like MerR and TetR.^28–30^ In addition, to our knowledge, our best design represents the most sensitive (low detection limit) CYA biosensor reported in the literature. More broadly, this work uncovers design rules for both *E. coli* whole-cell and cell-free biosensors composed of non-native transcriptional regulators. We anticipate that this work will have significant impact on the growing relevance of synthetic biology for rapid, point-of-use molecular detection.

## RESULTS

### Rational design of AtzR promoter variants in *E. coli* cells

To design a synthetic CYA biosensor functional in *E. coli*, we first considered the context of the natural CYA sensor. In *Pseudomonas* sp. strain ADP1, a divergent promoter comprising P*atzR* and P*atzDEF* transcribes *atzR* and the catabolic operon *atzDEF*, respectively, which are both regulated by AtzR.^24^ Regulation is coordinated through five AtzR binding sites in the intergenic region of P*atzDEF* and P*atzR*: two repressor binding sites (RBS-1 and RBS-2) and three activator binding sites (ABS-1, ABS-2 and ABS-3) (**Fig. 1a**).^24^ Unliganded, AtzR is a tetramer (dimer-of-dimers) where one dimer binds to RBS-1 and RBS-2 and the other dimer to ABS-2 and ABS-3. In this condition, transcription is repressed due to a bend in the DNA masking the σ^70^ −35 and −10 sites located on either side of ABS-3. Upon binding of CYA by AtzR, the dimers undergo a conformational change that relaxes their configuration on the DNA and causes one of the dimers to instead bind ABS-1 and ABS-2 (**Fig. 1b**).^31,32^ This “sliding dimer” mechanism results in exposure of the σ^70^ sites flanking ABS-3 and transcription of *atzDEF*.

**Figure 1.**
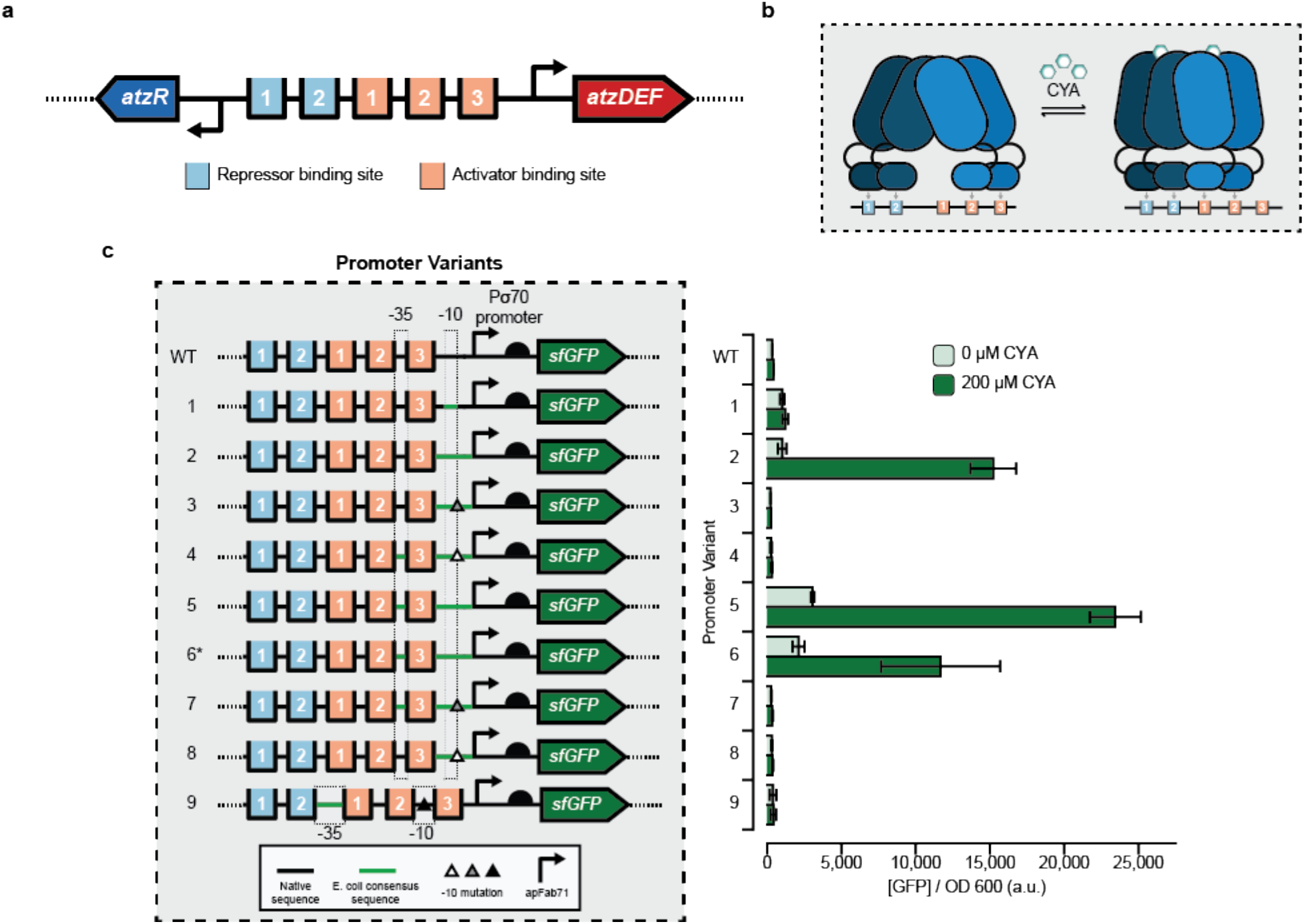
Engineering hybrid P*atz* operator-promoter variants for a cyanuric acid-inducible wholecell *E. coli* sensor. **(a)** The divergent P*atzDEF*-P*atzR* operon regulated by AtzR includes 2 repressor binding sites (light blue) that negatively autoregulate *atzR* expression and 3 activator binding sites (orange) that regulate *atzDEF* transcription. **(b)** AtzR, a LysR-Type Transcriptional Regulator (LTTR), is composed of a dimer-of-dimers. In the absence of cyanuric acid (CYA), AtzR binds to repressor binding sites 1 and 2, and activator binding sites 2 and 3. In the presence of CYA, a dimer of AtzR preferentially occupies activator binding sites 1 and 2, thereby allowing transcriptional machinery to access the promoter. This “sliding dimer” mechanism is typical of LTTRs. **(c)** Hybrid promoter designs to create a whole-cell *E. coli* sensor for CYA by combining native sequence from *Pseudomonas* sp. strain ADP1 at the 5’ end, a strong *E. coli* promoter (apFab71) at the 3' end and mutations to the transcriptionally-sensitive 3’ end of the −10 sequence (triangles). Transcriptional activation in the absence and presence of 200uM CYA was monitored by production of a downstream superfolder green fluorescent protein (sfGFP). A high-copy plasmid (a variant of the SC101origin of replication) was used for all promoter variants except promoter 6* which used a low-copy plasmid (p15a origin of replication). Bars indicate the average of experimental triplicate with error bars depicting 1 standard deviation.

To activate CYA-dependent transcription in *E. coli*, our strategy was to create hybrids of P*atzDEF* and a constitutive *E. coli* promoter (apFab71) without disrupting the arrangement of the RBS and ABS sites found in the native promoter. We hypothesized that a constitutive *E. coli* promoter would be necessary to provide transcriptional compatibility with the native polymerase of the whole-cell *E. coli* sensor, and that the TF activity could be maintained if the operator sites were kept with their appropriate spacing. The hybrid P*atzDEF*-apFab71 promoters were designed with AtzR binding sites of P*atzDEF* in the 5’ region of the promoter and apFab71 in the 3’ region (**Fig. 1c**). This arrangement allowed us to maintain the relative −35 and −10 positions in the hybrid promoters comparable to the native *Pseudomonas* promoter. We also included a well-characterized, *E. coli*-compatible 5’ UTR, including a ribosome binding site^33^ to improve translation initiation, and placed the entire construct upstream of an sfGFP reporter construct on a plasmid. AtzR was expressed under constitutive control from a second high-copy plasmid and the entire design was transformed into *E. coli* and characterized for activation in response to either 0 μM or 200 μM CYA supplemented to the media (**Fig. 1c**).

In total, we tested eight promoter variants with different sequences within the −35 and −10 sites in an attempt to vary transcriptional activity. The variants included combinations of the native P*atzDEF Pseudomonas* −35 site and σ^70^ consensus *E. coli* −10 site, σ^70^ consensus *E. coli* −35 and −10 sites, and single base substitutions at the transcriptionally-sensitive 3’ end of the σ^70^ consensus *E. coli* −10 site. In addition, one promoter design (P5) was tested on a low-copy (p15a origin of replication) plasmid (P6*) to assess whether lowering plasmid copy number would reduce sensor leak. Of these eight promoter variants, we identified three (P2, P5, and P6*) that gave strong transcriptional activation with CYA with fold-induction ratios of 15, 8 and 6, respectively.

Several design rules for LTTR-responsive promoters emerged from these results. First, *PatzR, PatzDEF* or the intragenic region between them gave very low reporter expression in both induced and uninduced states (**Supp. Fig. 1**). This result is likely due to weak crosstalk between the *E. coli* RNAP and the *Pseudomonas* σ^70^ recognition sites. Furthermore, even a strong ribosome binding site with a native *Pseudomonas* promoter gave weak reporter expression suggesting transcriptional incompatibility as the key bottleneck and a possible reason for poor signal in the previously published CYA sensor^27^. Second, hybrid *E. coli*-*Pseudomonas* promoters gave strong reporter expression. Among functional designs, promoters P5 and P6* are identical designs, but with the reporter expressed from either a high (P5) or low (P6*) copy number plasmid. Moving the reporter to a lower-copy plasmid left the OFF state largely unaffected, suggesting that availability of AtzR may be a stronger factor dictating the sensor’s OFF state than the reporter plasmid concentration. Because lowering the copy number decreased the sensor’s ON state by a factor of 2, the reporter plasmid copy number is likely the limiting factor when sensor is induced.

Though promoters P2, P3 and P4 are nearly identical, only promoter P2 was active. All three of these promoters had a *Pseudomonas* −35 site and the σ^70^ consensus *E. coli* −10 site, while promoters P3 and P4 each had a mutation at the 3’ end of −10 site. Similarly, P5, P7, and P8 all had *E. coli* σ^70^ consensus −35 and −10 sites and only P5 was functional; P7 and P8 also included single point mutations at the 3’ end of the −10 site. Taken together, our results suggest that an *E. coli* σ^70^ consensus −10 site is necessary but not sufficient for cyanuric acid-responsive transcription in *E. coli*. The addition of sequence elements from apFab71 in the 5’ UTR improved the transcriptional activity of native *Pseudomonas* promoter in *E. coli*.

### Dose-response and kinetics of the AtzR biosensor in *E. coli*

We chose P*atzDEF*-apFab71 promoter P5 for further characterization as it gave the highest sensor ON state. To comprehensively characterize our whole-cell biosensor, we next estimated its fold-induction ratio in triplicate at 8 different seeded culture densities, 12 different measurement times after induction, and 12 different concentrations of CYA (**Fig. 2a**). These parameters represent variables that must be optimized and tightly controlled for field testing applications using a whole-cell biosensor. Overall, we generated 384 total dose-response curves from 13,824 data points (**Supp. Fig. 2**).

**Figure 2.**
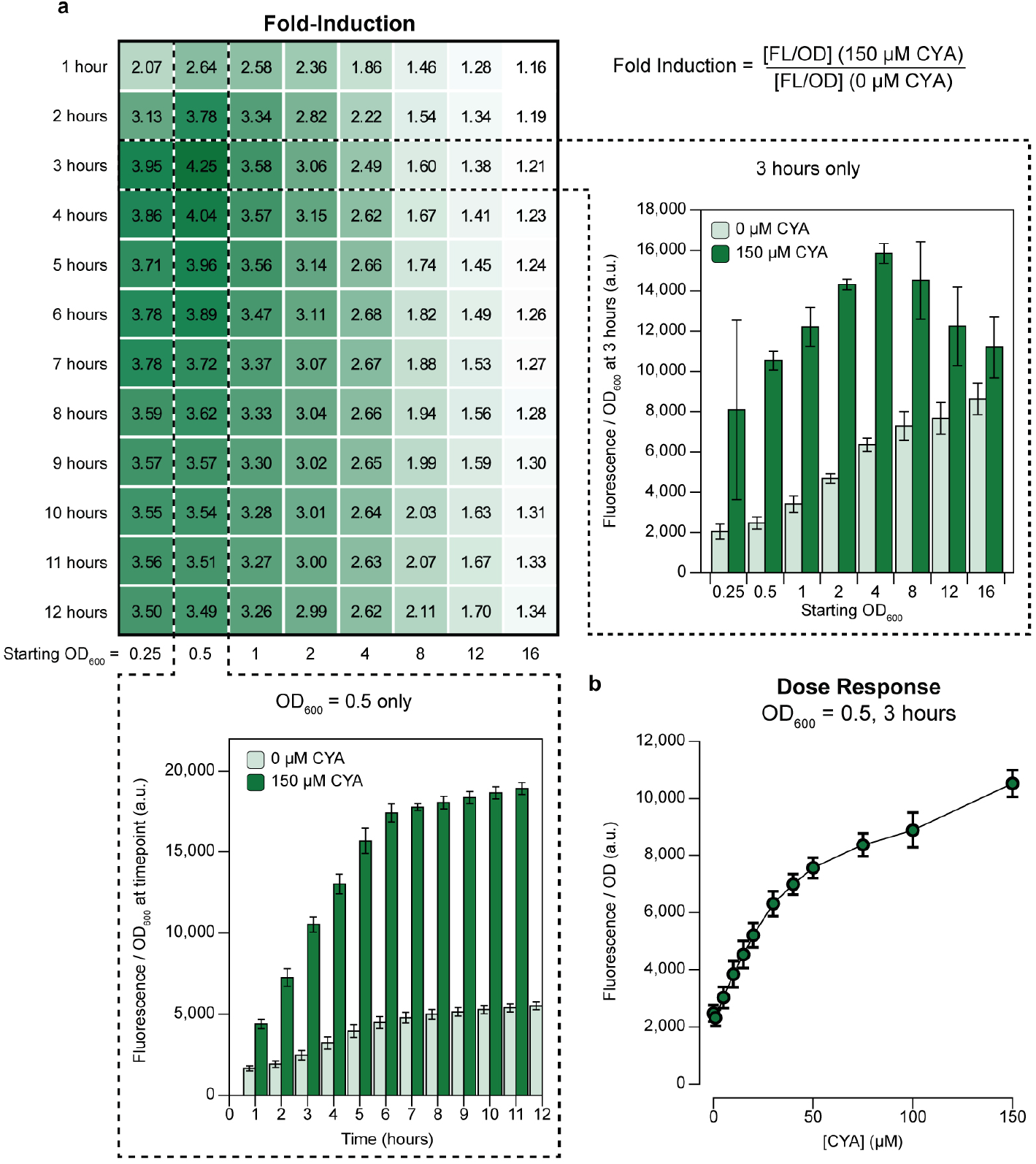
Performance of a whole-cell *E. coli* cyanuric acid sensor across time, optical densities, and CYA concentrations. **(a)** Fold-induction of the whole-cell cyanuric acid (CYA) sensor measured as ratio of reporter fluorescence in the presence or absence of 150 μM CYA. The starting cell optical densities (OD_600_) ranged from 0.25 to 16. Timepoints were taken every hour for 12 hours post-induction with CYA, and fold induction reported at all time points. The OD_600_ normalized sfGFP fluorescence time course for starting OD_600_ 0.5 is plotted with and without 150μM CYA (bottom). The OD_600_ normalized sfGFP fluorescence for starting OD_600_ at 0.5 to 16 at the 3-hour timepoint is plotted with and without 150 μM CYA (right). **(b)** Dose-response curve of the sensor measured 3 hours post-induction for starting OD_600_ 0.5. The OD_600_ normalized GFP fluorescence was taken from 0 to 150 μM CYA. All measurements were taken on 3 biological replicates. Averages are shown with error bars representing 1 standard deviation.

Our results reveal a smooth landscape for the sensor’s fold-induction, with an optimum fold-induction of ~4.25, which occurred at a relatively low seed inoculum (OD_600_ 0.5) and relatively early time step (3 hours). As both the time for measurement and the seeded optical density increase, the OFF state of the sensor increases more than the ON state (**Fig. 2a**, insets). After 1 hour of culturing, the sensor’s fold-activation is insensitive to time and virtually no change is observed beyond 6 hours of culturing. To evaluate the final analyte sensitivity of the biosensor, we measured reporter fluorescence over a wide range of CYA concentrations between 0 and 1 mM, supplied exogenously, at the optimal sensor conditions (optical density = 0.5; measurement time = 3 hours; **Fig. 2b and Supp. Fig. 2**). The sensor’s dose response increases monotonically with CYA concentration across 3 orders of magnitude, with a linear range including the recommended lower (25 ppm = 200 μM) and upper (75 ppm = 600 μM) concentrations of CYA in a swimming pool (**Supp. Fig. 2**).

In summary, we found the selected promoter (P5) has favorable characteristics, including low limit of detection, operational over a broad range of CYA concentrations, high signal-to-noise discrimination, and a moderately fast response time. However, our results also highlight the difficulties of using whole-cell biosensors for field-applications where sensitive instrumentation, time, and tight control over parameters do not exist. For example, although we found an optimal starting cellular density (OD_600_ = 0.5) and time-to-measurement (3 hours), a usable sensor would require costly instrumentation to measure both fluorescence and optical density, and adjust for these parameters as the cells continue to grow and divide. Furthermore, the population of cells comprising the sensor would have to remain relatively static from production to field-deployment.

### Design of a cell-free cyanuric acid sensor

With an eye towards developing a portable, on-demand platform that avoids the constraints of cellular biosensors, we next aimed to test whether cyanuric acid sensors identified in whole-cells can function in a cell-free extract (CFE) reaction. Although CFE systems have been developed from *Pseudomonas*^34^ (the native host of AtzR), *Pseudomonas* extracts produce far less protein than corresponding extracts prepared from *E. coli*, which have been optimized over several decades.^35–37^ Our success in developing a functional *E. coli* CYA sensor using the heterologous transcription factor AtzR thus motivated our approach to use *E. coli* extracts to build a cell-free sensor where AtzR regulates expression of sfGFP. We therefore designed two new constructs for cell-free expression: an AtzR-expressing plasmid and a reporter plasmid where sfGFP synthesis is inducible by CYA (**Fig. 3a**). By cloning the transcription factor and reporter into separate plasmids, the ratio of expression of each construct in the cell-free reaction can be tuned simply by changing the amount of each plasmid supplied to the reaction. This approach is simpler than the corresponding workflow for whole-cell biosensors, in which plasmid copy numbers must be manipulated while maintaining compatibility between origins of replication and selection markers.

**Figure 3.**
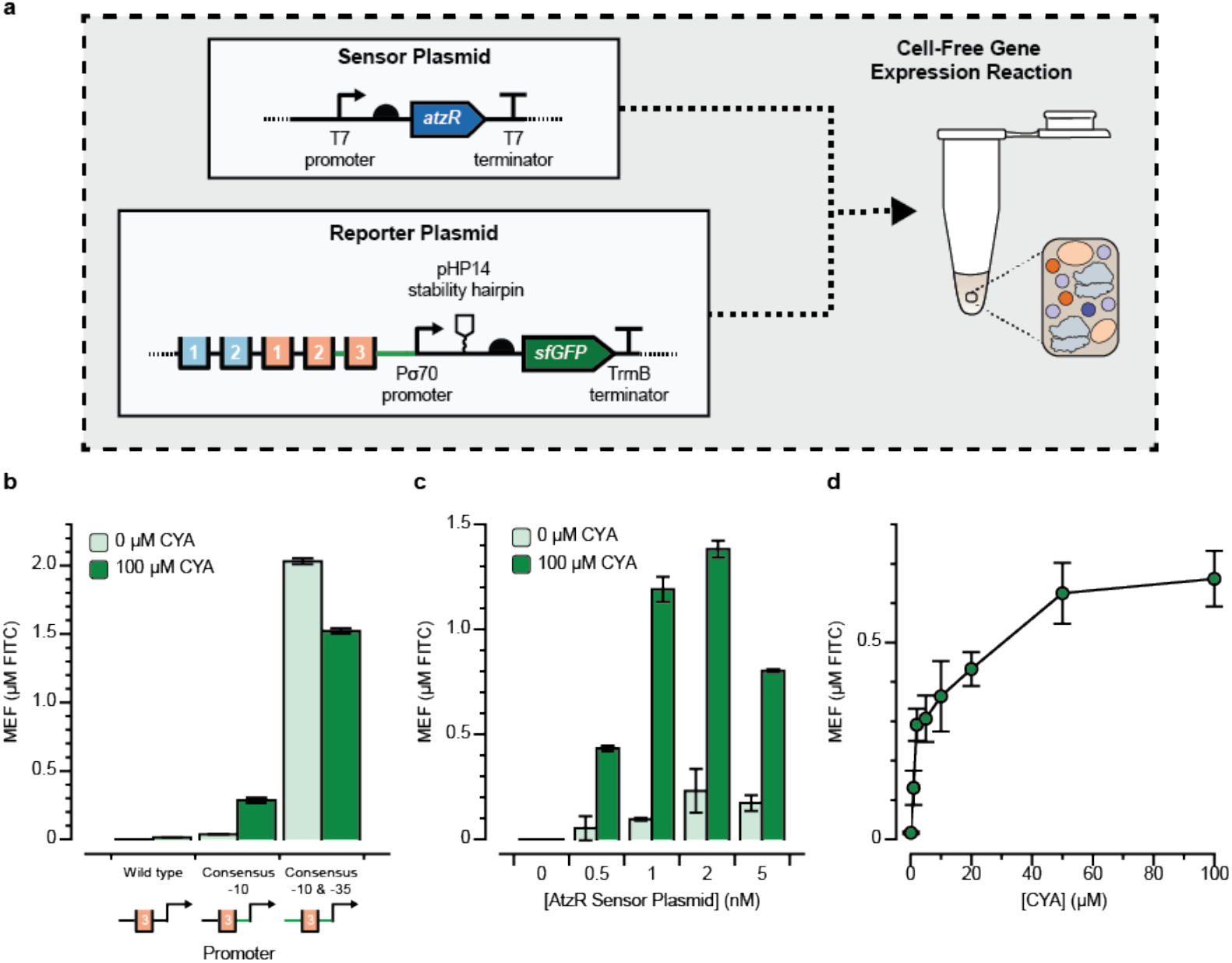
Design of a cell-free cyanuric acid sensor. **(a)** Two plasmids—one encoding the AtzR transcription factor (sensor plasmid) and one encoding sfGFP regulated by AtzR (reporter plasmid)—were added to 10 μL cell-free gene expression reactions and sfGFP production after four hours at 30°C was measured in the absence (OFF) and presence (ON) of 100 μM cyanuric acid (CYA). Plasmid designs are annotated as in **Fig. 1**. **(b)** Three promoter variants that replaced the native *Pseudomonas* σ^70^ recognition sites with the consensus *E. coli* −10 and −35 sites were tested in a small screen to identify promoter variants that were functional for cell-free CYA-induced activation. A promoter where only the −10 site was converted to the *E. coli* consensus sequence (TATAAT) gave the highest fold-induction and was carried through for subsequence experiments. In every case, reporter DNA plasmid was added at 10 nM, and the AtzR-expressing plasmid was kept at 1 nM. **(c)** Titration of AtzR-expressing plasmid against 10 nM reporter plasmid reveals that the optimal condition has 1 nM of AtzR-expressing plasmid to minimize sensor leak and potentially to maintain adequate distribution of translational resources between the two proteins. **(d)** Dose-response curve for cyanuric acid induction at optimal plasmid concentrations shows a wide operating range over 0-100 μM CYA. Kinetic plots showing the fluorescence trajectories for all samples in b-d are in **Supp. Fig. 3**. Error bars represent 1 standard deviation from 3 technical replicates. Fluorescence is reported using mean equivalent fluorescence (MEF) of fluorescein isothiocyanate based on a previously developed standard curve.

To design cell-free expression constructs, we rationally adjusted the constructs that were functional *in vivo* to better resemble cell-free expression cassettes. In the AtzR sensor plasmid, we replaced the constitutive bacterial promoter to a T7 promoter, since T7 RNAP is the most common and productive polymerase for making proteins in cell-free expression systems, resulting in an enhancement of protein yield by a factor of 2-3.^38,39^ For the reporter plasmid, we started with a design based on Promoter P5, since it gave both a high ON signal and fold-induction *in vivo*, and previous work has shown that very strong promoters are necessary for robust protein synthesis in extract.^39^ We additionally included a pHP14 RNA stability hairpin immediately after the transcriptional start site in the reporter plasmid, since secondary structure in the mRNA 5’ end is necessary for robust expression from bacterial promoters in cell-free conditions.^39^ Both expression cassettes were then placed into high-copy vectors.

The two plasmids were then purified and doped into a cell-free reaction containing postlysis processed *E. coli* extract^39^ and the additional biological cofactors required for transcription and translation *in vitro*. The reactions were incubated at 30°C for 4 hours and the reaction progress was monitored by continuously measuring fluorescence on a plate reader. sfGFP fluorescence from these reactions was then standardized to a NIST-traceable solution of fluorescein isothiocyanate (FITC), to facilitate comparison across experiments (see Methods).^39^

We observed constitutive transcription in our initial reporter design based on Promoter P5 (with consensus *E. coli* σ^70^ −10 and −35 sites) even in the absence of CYA (**Fig. 3b**). This result was unexpected as it contradicts the results obtained in cells. We hypothesized that variability in reaction environments between cells and in CFE, particularly the concentration of the accessory factors for transcription, gave rise to this difference. We therefore prototyped new reporter designs and observed that, surprisingly, a promoter based on Promoter P2 with the wild-type *Pseudomonas* −35 sequence but the consensus *E. coli* σ^70^ −10 sequence, responded well to 100 μM cyanuric acid *in vitro* (**Fig. 3b**) with ~8-fold-induction and low sensor leak. We thus applied this design to further optimize the cell-free sensor.

We next aimed to find the optimal ratio between the AtzR-expressing plasmid and the reporter plasmid that gives the best fold-induction. We therefore fixed the sfGFP reporter plasmid concentration in the reaction to 10 nM and varied the concentration of the AtzR-expressing plasmid. Fluorescence measurements were taken after a 4-hour incubation in either the absence (OFF) or presence (ON) of 100 μM CYA (**Fig. 3c**). When no AtzR-expressing plasmid was supplied to the CFE reaction, we observed no sfGFP production, consistent with an activation mechanism. As the AtzR template concentration increased, we observed an increase in both the ON and OFF states, up to 2 nM AtzR plasmid. Beyond 2 nM of the AtzR plasmid, sfGFP production tapered off, likely due to competition for translational resources between reporter and transcription factor synthesis in the batch reaction. We chose 1 nM AtzR plasmid and 10 nM reporter plasmid as the optimal condition, which achieves >12-fold-induction in the presence of 100 μM CYA. These results highlight the power of cell-free systems for rapidly tuning sensors, since the analogous cellular experiments would have required extensive cloning and testing.

Next, we measured the cell-free sensor’s dose response against CYA. At the optimal conditions (10 nM reporter plasmid, 1 nM AtzR-expressing plasmid), we observed a wide operating range of the sensor between 1 and 50 μM CYA, with maximal activation achieved at 100 μM (**Fig. 3d**). The cell-free optimized dose-response matches the in-cell dose-response curve both quantitatively and qualitatively, and our data suggest that the sensor can detect the EPA’s recommended lower and upper limit for CYA in swimming pools (200 and 600 μM) if diluted into a prepared reaction at a 1:10 ratio.

### Proof-of-principle cell-free cyanuric acid detection at the point-of-use

With a cell-free CYA sensor at hand, we aimed to test if our cell-free sensor could directly detect CYA in natural and synthetic water samples, as a final validation that a cell-free biosensor can be deployed for reporting on relevant concentrations of CYA outside the laboratory. First, we validated that CYA does not inhibit cell-free transcription and translation at concentrations below 10 mM, which is far above the highest expected concentration of CYA in pool water (600 μM) (**Supp. Fig. 4**). Next, we freeze-dried premixed reactions containing the sensor and reporter plasmids, reaction buffer, and extract, and rehydrated them with water samples obtained from a drinking fountain, Lake Michigan (Evanston, IL), and a chlorinated local swimming pool (**Fig. 4b**). In each case, we spiked one reaction with 100 μM CYA to serve as a control for the impact of water quality on the cell-free reaction’s performance. Importantly, to better reflect real-world operating conditions for such a sensor, we did not filter or otherwise process any of the water samples. We then measured sfGFP production on a plate reader over 4 hours.

**Figure 4.**
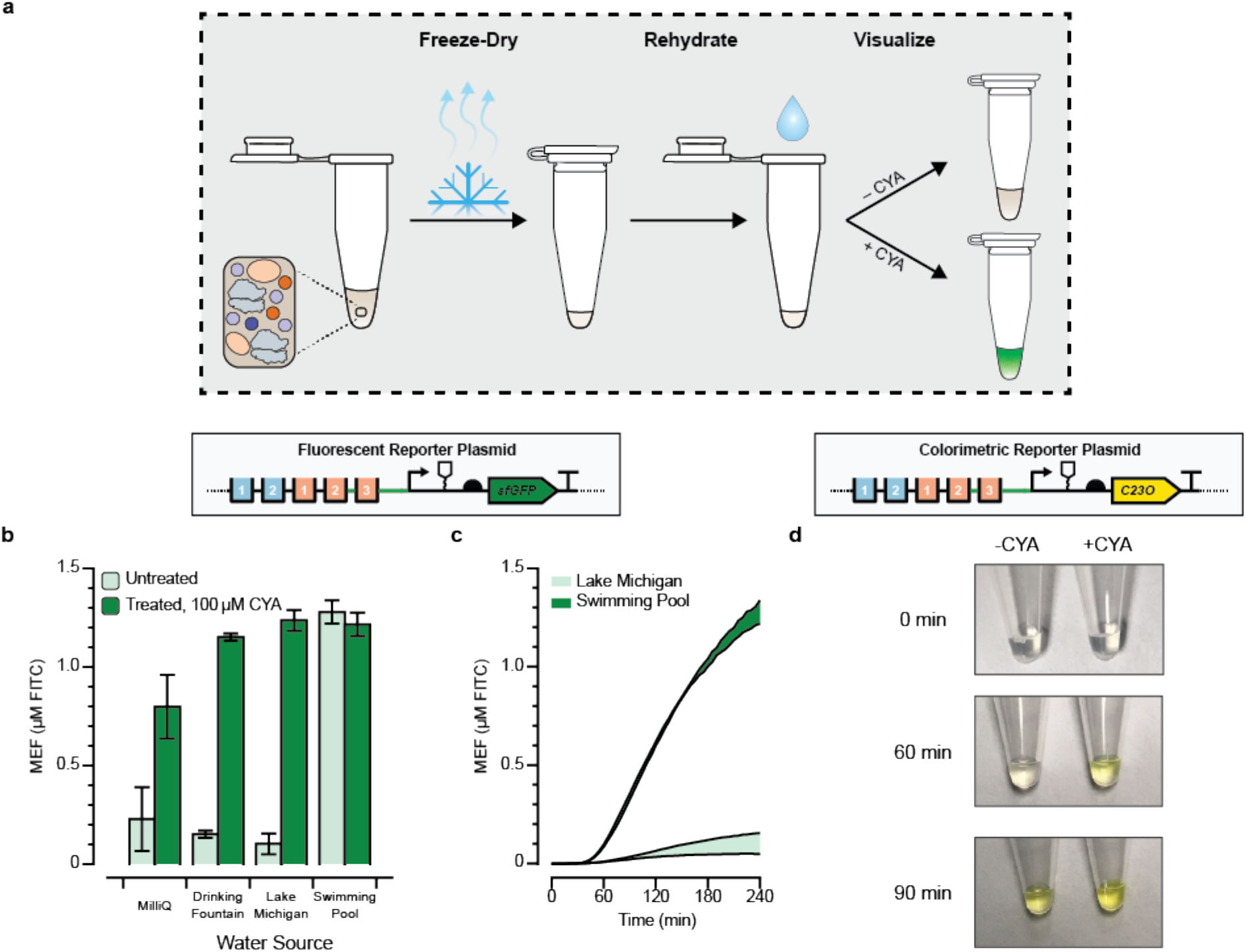
Proof-of-concept for field-deployable cyanuric acid sensing using a cell-free biosensor. **(a)** Optimized cell-free reactions are prepared without cyanuric acid (CYA), freeze-dried, and then rehydrated with unfiltered environmental water samples. **(b)** CYA in environmental water samples can be detected from freeze-dried cell-free reactions. Pre-mixed cell-free reactions were freeze-dried and rehydrated with 90% of an environmental water sample and 10% of either MQ water (OFF) or 10% MQ water spiked with a final concentration of 100 μM CYA (ON). sfGFP fluorescence is specifically induced by CYA in every sample except the pool water, which is expected to already contain CYA even in the OFF state. The kinetics for all conditions is presented in **Supp. Fig. 5**. **(c)** Time course comparisons of the OFF state (No CYA added) from panel B, for the unfiltered environmental water samples that were not spiked with CYA. Pool water can be distinguished against lake water in less than one hour. **(d)** Time course comparison for activation of cell-free reaction driving expression of catechol 2,3-dioxygenase, an enzymatic reporter that cleaves its colorless substrate catechol into a yellow product, *cis,cis*-muconic acid. CYA was added at a concentration of 600 μM in the ON state to a fresh cell-free reaction incubated at 30°C and monitored for 90 minutes. Pictures are cropped representative images from the experiment, which was done in triplicate. Uncropped images are in **Supp. Fig. 7**. Error bars in **b** and **c** represent one standard deviation from three technical replicates. Fluorescence in **b** and **c** is reported standardized to mean equivalent fluorescence (MEF) of fluorescein isothiocyanate based on a previously developed standard.

For the samples hydrated with laboratory water, fountain water, and lake water, the ON states created by spiking in CYA showed appreciable activation versus the OFF states with no CYA added, and similar sfGFP yields were obtained to the yields from fresh reactions in the earlier experiments. This result shows that the reactions are not poisoned by the unfiltered environmental water samples that we tested (**Fig. 4b, Supp. Fig. 5**). The pool water sample, which was expected to have CYA, was activated even without the CYA spike-in (**Fig. 4b**) and showed activation relative to the lake water sample in less than an hour (**Fig. 4c**). We conclude that our cell-free biosensor is able to detect whether or not an arbitrary environmental water sample contains CYA. Since we did not observe sample matrix poisoning effects in this experiment, we hypothesize that a standard curve of CYA samples could thus be developed in the future as a method for quantification of CYA concentrations at the point-of-use.

Finally, we aimed to validate if a visible output signal for the sensor could be generated without a plate reader to enable equipment-free detection of CYA from environmental samples. Specifically, we prototyped a sensor to report water samples with dissolved CYA above the upper recommended concentration of 600 μM. Because fluorescence of sfGFP is difficult to detect by the unaided eye, particularly in the early stages of the reaction, we replaced the coding sequence of the reporter plasmid instead with the enzyme catechol 2,3-dioxygenase (C23DO). C23DO cleaves its substrate, catechol, into a yellow *cis,cis-* muconic acid product, so the reaction progress can be directly read out from the visible color.^40^ By decreasing the concentration of the C23DO reporter plasmid in the reaction to 5 nM, we observed robust activation of the sensor in the presence of 600 μM CYA and no leak within 2 hours, when read out on a plate reader (**Supp. Fig. 6**).

We then prepared the same experiment in tubes maintained at 30°C and visually monitored the reaction progress over time. Activation of the reporter was observed by the production of a yellow color within 1 hour (**Fig. 4d**; **Supp. Fig. 7**). Interestingly, when the sensing reaction was prepared in tubes instead of the plate reader, leak was observed on longer timescales, such that the OFF state was fully activated within 90 minutes. Our results here hint that further optimization of the reaction conditions will be necessary to make a colorimetric reporter viable for quantification of cyanuric acid in a sample. Nonetheless, this experiment provides a powerful proof-of-principle for equipment-free, field-deployable detection of cyanuric acid at relevant concentrations.

## DISCUSSION

In this work, we demonstrate a strategy for designing field-deployable biosensors by using non-native LTTR transcriptional regulators from *Pseudomonas* sp. strain ADP1in *E. coli* cells and cell-free extracts. This strategy had several key features. First, sequence-to-function data was enabled by modifying the DNA-binding sequence (promoter) for the metabolite-responsive transcription factor AtzR using a hybrid promoter library composed of native (*e.g., Pseudomonas*) and synthetic (*e.g., E. coli*) host components. We have shown that the AtzR-activated native promoter from *Pseudomonas sp*. strain *ADP1* suffers from a low expression and activation levels in *E. coli*. The native promoter with a strong *E. coli* ribosome binding site also gave low expression suggesting weak transcriptional compatibility in the non-native host or possible presence of inhibitory sequence elements. On the other hand, our rationally designed hybrid promoters from a strong *E. coli* promoter and the native promoter without disrupting native binding sites gave much higher expression and ligand-responsive activation in *E. coli*. Few LTTR sensors have previously been reported in the literature, and we anticipate that this work will shed light on the design rules necessary to engineer more sophisticated biosensors for this very large class of natural allosteric transcription factors.

Second, we demonstrated that functional sensor designs identified in whole-cells could be used to inform the development of cell-free biosensors. This is important because it set the stage to use cell-free systems to avoid several practical limitations that arise in whole-cell biosensors, including the fact that mechanisms in cells have evolved to facilitate species survival that may compromise sensor robustness, or mutate the biosensing machinery. This result invites questions on the agreement between whole-cell and cell-free biosensors. After minor optimizations, we observed similar dose-response curves between the two sensors, with each attaining around 5-10 fold-activation in response to similar concentrations of cyanuric acid. Interestingly, the cell-free sensor appears to hit its saturation regime at lower concentrations than the cellular sensor. This could be consistent with a possible mechanism where transport of CYA through the cell membrane is rate-limiting. Another key difference occurs in the optimal promoter design. In cells, the best-fold activation is achieved when the *E. coli* consensus sequence is used at both σ^70^ recognition sites, but this promoter was constitutively ON when tested in cell-free conditions. We hypothesize that this difference may be attributable to differing concentrations of the *E. coli* RNAP *holo* complex and the corresponding sigma factors in cells versus *in vitro*. Indeed, previous work has shown that some quorum sensing systems show appreciable differences in behavior when ported between cells and cell-free conditions.^41^ Thus, it is evident that each system’s parameters (e.g., plasmid copy numbers for whole-cell sensors) must be optimized in their own specific context.

Third, we showed that our freeze-dried, cell-free system can detect cyanuric acid at environmentally relevant concentrations in natural water samples within a single hour, incubated at or below body temperature. This result is important not just for controlling chlorine levels in swimming pools, but also potentially for measuring the concentration of more toxic pollutants in runoff streams, since cyanuric acid is a natural catabolite of the widely overused herbicide atrazine. Toward this end, we are encouraged by the fact that our field-deployable platform is the most sensitive reported CYA biosensor in the literature. It is also inexpensive. We estimate that the entire freeze-dried reaction, including the colorimetric substrate catechol, should only cost around $0.05/tube (**Supp. Table 1**).

Looking forward, our work offers promise for the deployment of a wide array of biosensors for detection of toxic molecules in the environment. However, future work should be done to tune the sensor’s response to a specific desired threshold concentration. The large operating range observed in both the cellular and cell-free sensors could only enable fluorescence quantification in the presence of a standard curve, which is not ideal for field-deployment. The use of a colorimetric enzymatic output, such as catechol 2,3-dioxygenase, highlights a path forward to address this issue. This work hints at an exciting future where cell-free molecular sensors freeze-dried and packaged in ambient conditions, effectively function as water quality diagnostics in field conditions at the point-of-need.

## MATERIALS AND METHODS

### Plasmid design and preparation

The sensor plasmid for the cellular sensor was constructed on a backbone carrying a spectinomycin resistance gene and SC101 origin of replication. It uses a constitutive promoter apFAB61 and a ribosome binding site BBa_J61132 (**Supp. Table 2**) to drive the expression of AtzR (codon optimized for *E.coli*). Reporter plasmids were constructed with a backbone carrying a kanamycin resistance gene and a ColE1 origin of replication. Super-folder green fluorescent protein (sfGFP)^42^ was used as the reporter gene with a strong ribosome binding site (Bujard).^33^ Promoters, as described in the results, were inserted into the plasmids to drive the expression of the reporter gene. All DNA fragments were amplified with a high-fidelity KAPA DNA polymerase (Kapa Bioscience KK2101) and assembled into plasmids using isothermal assembly.^43^ Assembled reactions were transformed into DH10B *E. coli* cells (New England Biolabs Inc, Cat. C3020), plated on LB-agar, and screened for kanamycin (50 μg/mL) resistance. Screened colonies were then subcultured with shaking at 37°C for 16 hours in LB. Plasmids were isolated from subcultures using a DNA miniprep kit following the manufacturer’s instructions (Omega BioTek) and were Sanger sequenced to identify correctly assembled plasmids.

The main reporter plasmid (JBL7030) for cell-free expression was constructed using inverse PCR and blunt-end ligation based on our previously reported plasmid pJBL7023,^39^ which encodes the strong constitutive promoter J23119, the PHP14 RNA stability hairpin, a strong ribosome binding site, and the coding sequence for sfGFP. The AtzR binding sequence was inserted immediately 5’ to the stability hairpin and the promoter was engineered by adapting the annotated −10 site to the *E. coli* σ70 consensus sequence TATAAT. The −35 site sequence was kept as CAGTCA. The C23DO reporter variants were designed from this starting plasmid by replacing the sfGFP coding sequence using isothermal assembly. The sensor plasmid (JBL7032) for cell-free expression was designed based in the pJL1 cell-free expression vector, where AtzR was expressed under control of a T7 promoter, and cloned using isothermal assembly.^43^ Relevant plasmid sequences are included in the Supplementary Table 2 All plasmids were purified for cell-free expression using Qiagen Plasmid Midiprep Kit (Cat. 12145) and quantified by Qubit dsDNA BR Assay Kit (Invitrogen #Q32853).

### *In vivo* fluorescence measurement

Reporter and sensor plasmids (including WT promoter plasmid) were cotransformed into DH10B cells (New England Biolabs Inc, Cat. C3020) using electroporation and then plated onto LB-agar plates with kanamycin (50 μg/mL) and spectinomycin (50 μg/mL). After overnight growth at 37°C, at least 3 individual colonies for each plasmid pair were picked and inoculated into 150 μL LB with the above-mentioned antibiotics in a 96-well plate. After 3 hours of growth at 37°C in a multi-plate shaker at 1100 rpm (Southwest Science, Cat. SBT1500-H), cells were diluted 1:50 into 150 μL MOPS medium (supplemented with 0.4% glucose and 1X BME vitamin) with antibiotics in a 96-well plate. Cyanuric acid, prepared in DMSO, was introduced to inducing wells at a final concentration of 200 μM, with DMSO at final volume of 1%. Non-inducing wells also received same final volume of DMSO. After 16 hours at 37°C with shaking (1100 rpm), optical density at 600nm (OD_600_) and GFP fluorescence (excitation at 488 nm and emission at 520 nm, gain of 40) of each well was measured using a multi-well plate reader (Synergy HTX, BioTek Inc). Fluorescence was normalized to cell density measured at OD_600_. A blank value (normalized fluorescence of cells without a GFP reporter) was subtracted from those measurements. Mean and standard deviation of each reporter plasmid activity were calculated from measurements of 3 biological replicates.

### Optimal conditions for *in vivo* application

To find optimal conditions for *in vivo* application of the sensor, we tested the effect of starting optical density at 600nm (OD_600_) and concentration of CYA. We inoculated fresh colonies of P5 with the sensor plasmid into 3 mL LB with 50 μg/mL of both kanamycin and spectinomycin. After overnight growth, the culture was inoculated into 300 mL LB with antibiotics. The cultures continued to grow at 37°C under shaking (225 rpm, 3 cm radius) until the OD_600_ reached 0.5. Cell cultures were then centrifuged at 4000 x g for 5 minutes. After removing the medium, the cell pellet was resuspended in 9.4mL warm fresh LB with antibiotics (final OD_600_ = 16). Following the resuspension, we diluted culture into other desired OD_600_ with warm fresh LB (with 50 μg/mL kanamycin and spectinomycin) and added 150 μL of culture into each well of a 96-well plate. CYA in DMSO was added to cells at appropriate inducer concentrations as described above. The plate was then immediately transferred to a multi-well plate reader (Synergy HTX, BioTek Inc) with continuous shaking at 37°C. The fluorescence of each well was recorded by the plate reader (excitation 448 nm, emission at 520 nm, gain of 40) every 15 minutes. Data was collected for 12.5 hours with 3 biological replicates.

### Cell-free extract preparation and protein synthesis

Cell-free extract was prepared following our published method^44^ using a BL21 (DE3) host strain genomically modified to lack endogenous beta-galactosidase (LacZ). Briefly, a 1 L culture of the host strain was prepared in 2xYT + P media and grown to OD_600_ = 3.0. Cells were harvested by centrifugation, lysed by sonication, and centrifuged to isolate the supernatant. An 80-minute ribosomal run-off reaction was performed on the supernatant at 37°C, followed by a 3-hour dialysis at 4°C. The LacZ deficient (lac operon deletion) BL21-Gold-dLac (DE3) strain was a gift from Jeff Hasty (Addgene plasmid # 99247).^29^

The cell-free protein synthesis reaction was run as previously described^39^, using 10 nM of the reporter plasmid and 1 nM of the sensor plasmid in all reactions after optimization. For sfGFP synthesis reactions, fluorescence was measured by excitation at 485 nanometers and emission at 520 nanometers in a BioTek Synergy H1 plate reader in a black 384 well-plate (Corning 3712). sfGFP fluorescence was correlated to concentration of FITC by a previously developed calibration standard.^44^ All reactions were run in triplicate at the 10 μL scale at 30°C for 4 hours. For the colorimetric assay, 2 mM catechol was supplemented to the reaction in addition to the C23DO reporter plasmid at 5 nM. These reactions were incubated at 30°C for 2 hours, with photos taken every 15 minutes using an iPhone with the iOS camera app (iPhone SE).

For the freeze-dried experiments, full reactions without cyanuric acid were assembled on ice in PCR tubes, snap-frozen on liquid nitrogen, and then lyophilized overnight on a Labconco FreeZone 2.5L Bench Top Freeze Dry System under 0.04 mbar pressure and −85°C condenser temperature. After approximately 16 hours of lyophilization, the reactions were rehydrated with the full volume of liquid that was dried off, incubated for 1 minute at 37°C to fully dissolve the reaction pellet, and then dispensed in triplicate into a well-plate. For sample matrix experiments, the rehydration was prepared with and without cyanuric acid by performing a 1:10 dilution of 1 mM stock of cyanuric acid dissolved in laboratory-grade water into lake, pool, or tap water, and then rehydrating the reaction with the dilutions.

## ACKNOWLEDGEMENTS

The authors would like to acknowledge members of the Raman, Jewett and Lucks Labs for helpful discussions. S.R. and X.L. were supported by the University of Wisconsin-Madison and Army Research Office grant W911NF1710043. M.C.J. acknowledges support from the Air Force Research Laboratory Center of Excellence Grant FA8650-15-2-5518, the DARPA 1000 Molecules Program HR0011-15-C-0084, the David and Lucile Packard Foundation, and the Camille Dreyfus Teacher-Scholar Program. J.B.L. acknowledges support from the Air Force Research Laboratory Center of Excellence Grant FA8650-15-2-5518, the Camille Dreyfus Teacher-Scholar Program and Searle Funds at the Chicago Community Trust. A.D.S. was supported in part by the National Institutes of Health Training Grant (T32GM008449) through Northwestern University’s Biotechnology Training Program. The U.S. Government is authorized to reproduce and distribute reprints for Governmental purposes notwithstanding any copyright notation thereon. The views and conclusions contained herein are those of the authors and should not be interpreted as necessarily representing the official policies or endorsements, either expressed or implied, of the Army Research Office, Air Force Research Laboratory, Air Force Office of Scientific Research, DARPA, or the U.S. Government.

