## Supplementary Information for "Design of a transcriptional biosensor for the portable, on-demand detection of cyanuric acid"

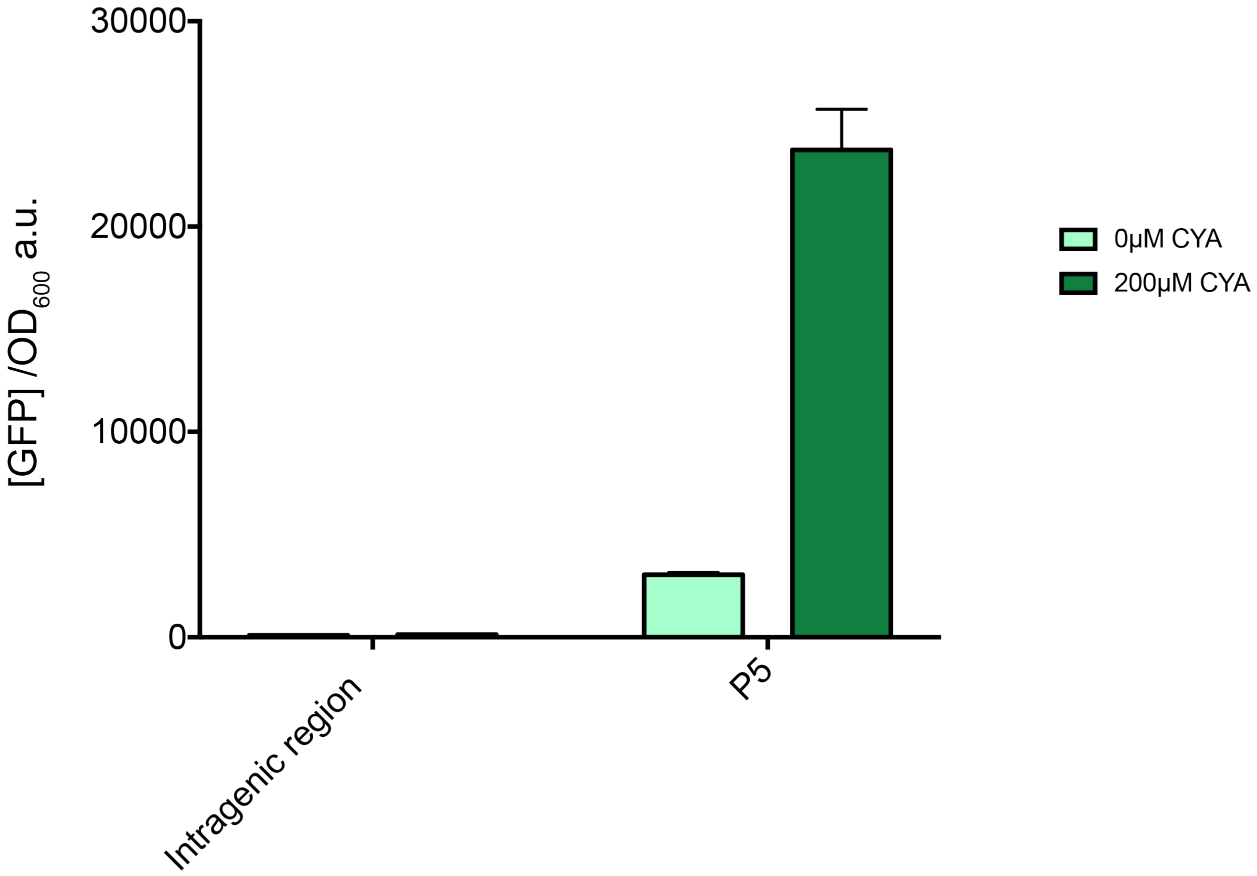


**Supplemental Figure 1: Cellular expression comparison between atzRDEF intragenic region and hybrid promoter P5.** Transcriptional activation was monitored by production of a downstream superfolder green fluorescent protein (sfGFP). Expression levels are calculated as sfGFP fluorescence (excitation: 488nm emission: 510nm) over cell optical density (at 600nm). Bars indicate the average of experimental triplicate with error bars depicting ± standard deviation.

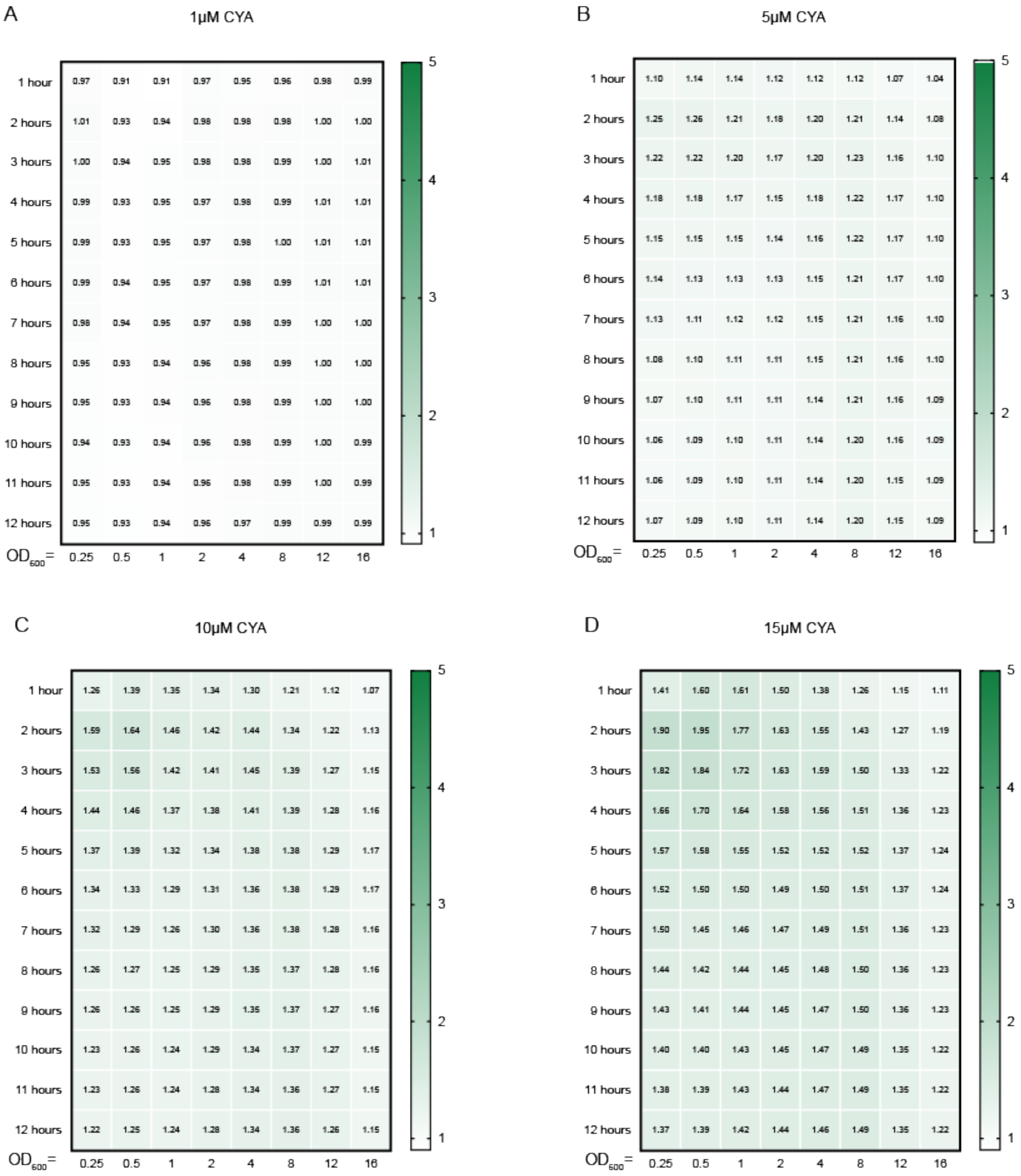


**Supplemental Figure 2**. **Performance of a whole-cell *E. coli* cyanuric acid sensor across time and optical densities at all tested CYA concentrations.** Fold-induction of the whole-cell cyanuric acid (CYA) sensor measured as ratio of reporter fluorescence in the presence or absence of 1 μM (**A**), 5 μM (**B**), 10 μM (**C**), 15 μM (**D**), 20 μM (**E**), 30 μM (**F**), 40 μM (**G**), 50 μM (**H**), 75 μM (**I**), 100 μM (**J**), 150 μM (**K**), 190 μM (**L**), 385 μM (**M**), 580 μM (**N**), 770 μM (**O**), 960 μM (**P**) CYA. Average fold-inductions of 3 biological replicates are shown.

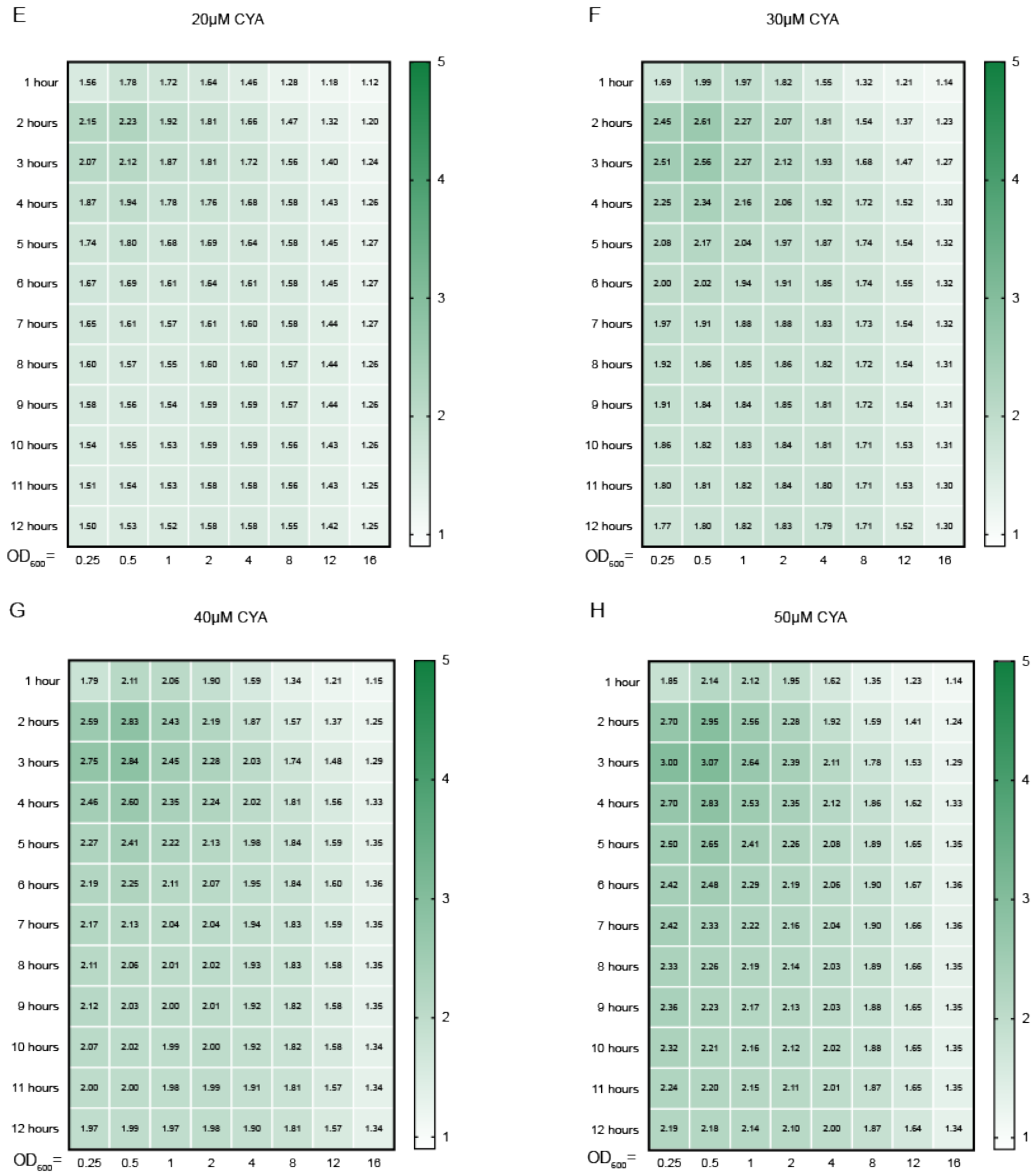


**Supplemental Figure 2**. **Performance of a whole-cell *E. coli* cyanuric acid sensor across time and optical densities at all tested CYA concentrations.** Fold-induction of the whole-cell cyanuric acid (CYA) sensor measured as ratio of reporter fluorescence in the presence or absence of 1 μM (**A**), 5 μM (**B**), 10 μM (**C**), 15 μM (**D**), 20 μM (**E**), 30 μM (**F**), 40 μM (**G**), 50 μM (**H**), 75 μM (**I**), 100 μM (**J**), 150 μM (**K**), 190 μM (**L**), 385 μM (**M**), 580 μM (**N**), 770 μM (**O**), 960 μM (**P**) CYA. Average fold-inductions of 3 biological replicates are shown.

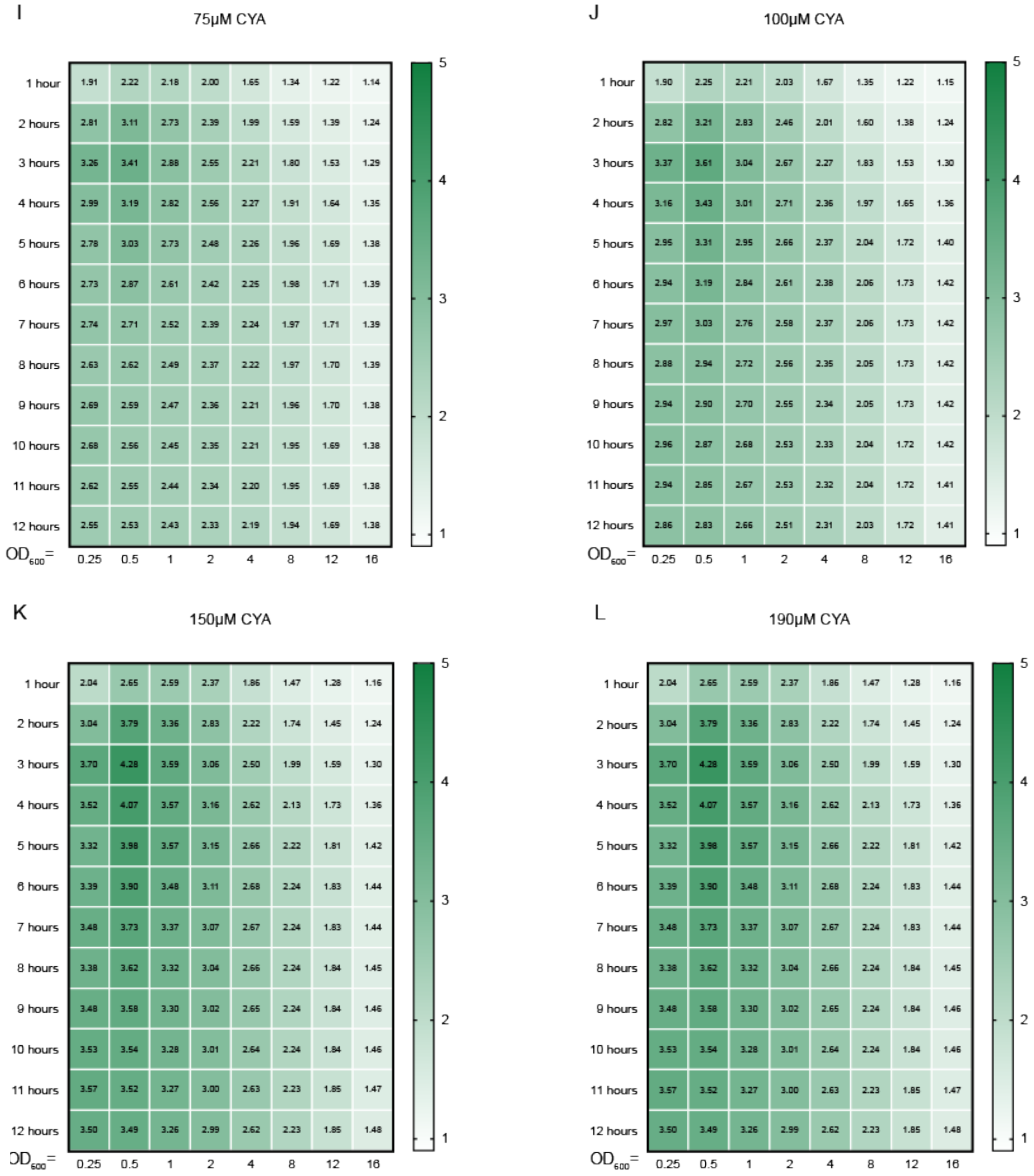


**Supplemental Figure 2**. **Performance of a whole-cell *E. coli* cyanuric acid sensor across time and optical densities at all tested CYA concentrations.** Fold-induction of the whole-cell cyanuric acid (CYA) sensor measured as ratio of reporter fluorescence in the presence or absence of 1 μM (**A**), 5 μM (**B**), 10 μM (**C**), 15 μM (**D**), 20 μM (**E**), 30 μM (**F**), 40 μM (**G**), 50 μM (**H**), 75 μM (**I**), 100 μM (**J**), 150 μM (**K**), 190 μM (**L**), 385 μM (**M**), 580 μM (**N**), 770 μM (**O**), 960 μM (**P**) CYA. Average fold-inductions of 3 biological replicates are shown.

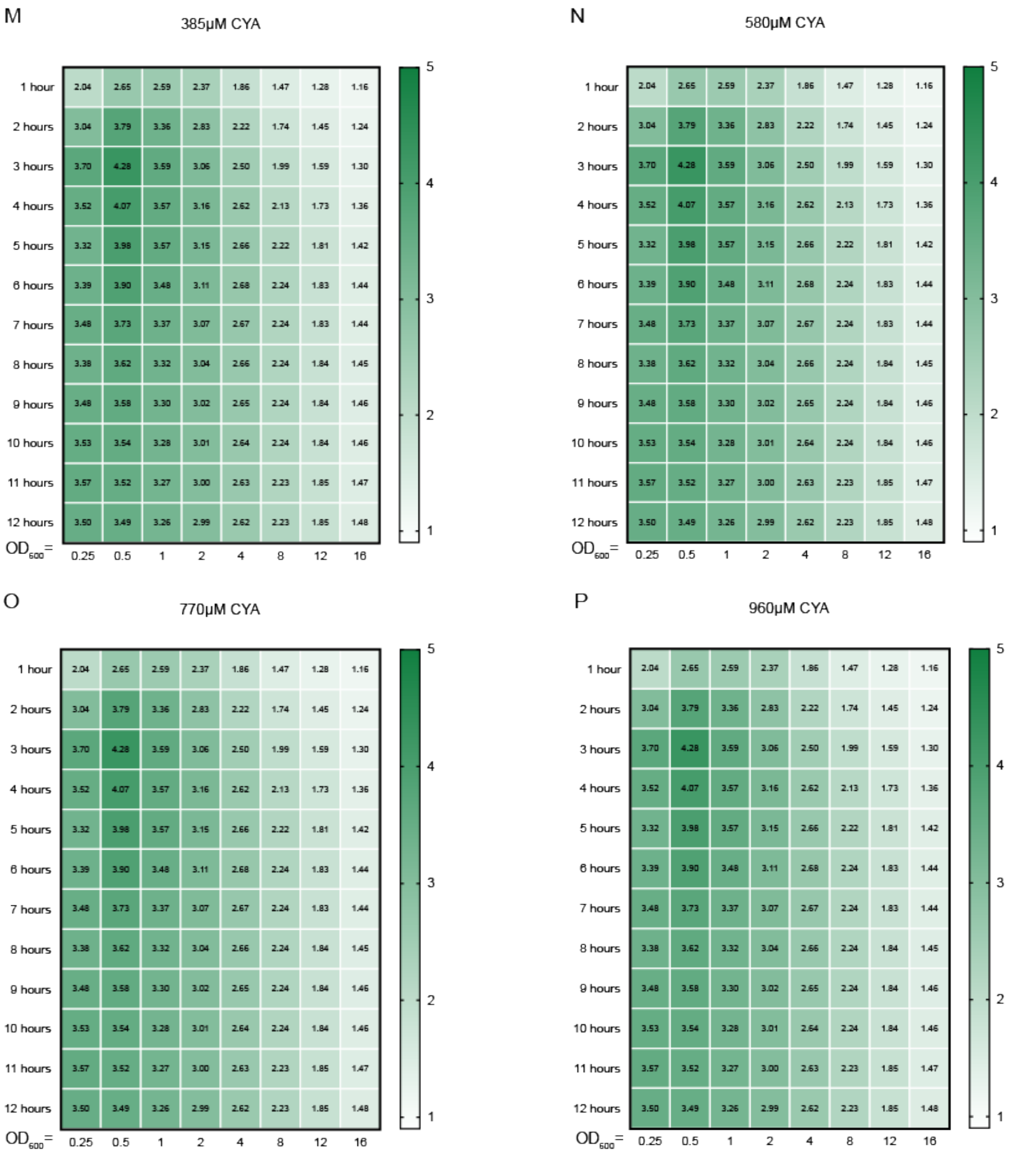


**Supplemental Figure 2**. **Performance of a whole-cell *E. coli* cyanuric acid sensor across time and optical densities at all tested CYA concentrations.** Fold-induction of the whole-cell cyanuric acid (CYA) sensor measured as ratio of reporter fluorescence in the presence or absence of 1 µM (**A**), 5 µM (**B**), 10 µM (**C**), 15 µM (**D**), 20 µM (**E**), 30 µM (**F**), 40 µM (**G**), 50 µM (**H**), 75 µM (**I**), 100 µM (**J**), 150 µM (**K**), 190 µM (**L**), 385 µM (**M**), 580 µM (**N**), 770 µM (**O**), 960 µM (**P**) CYA. Average fold-inductions of 3 biological replicates are shown.

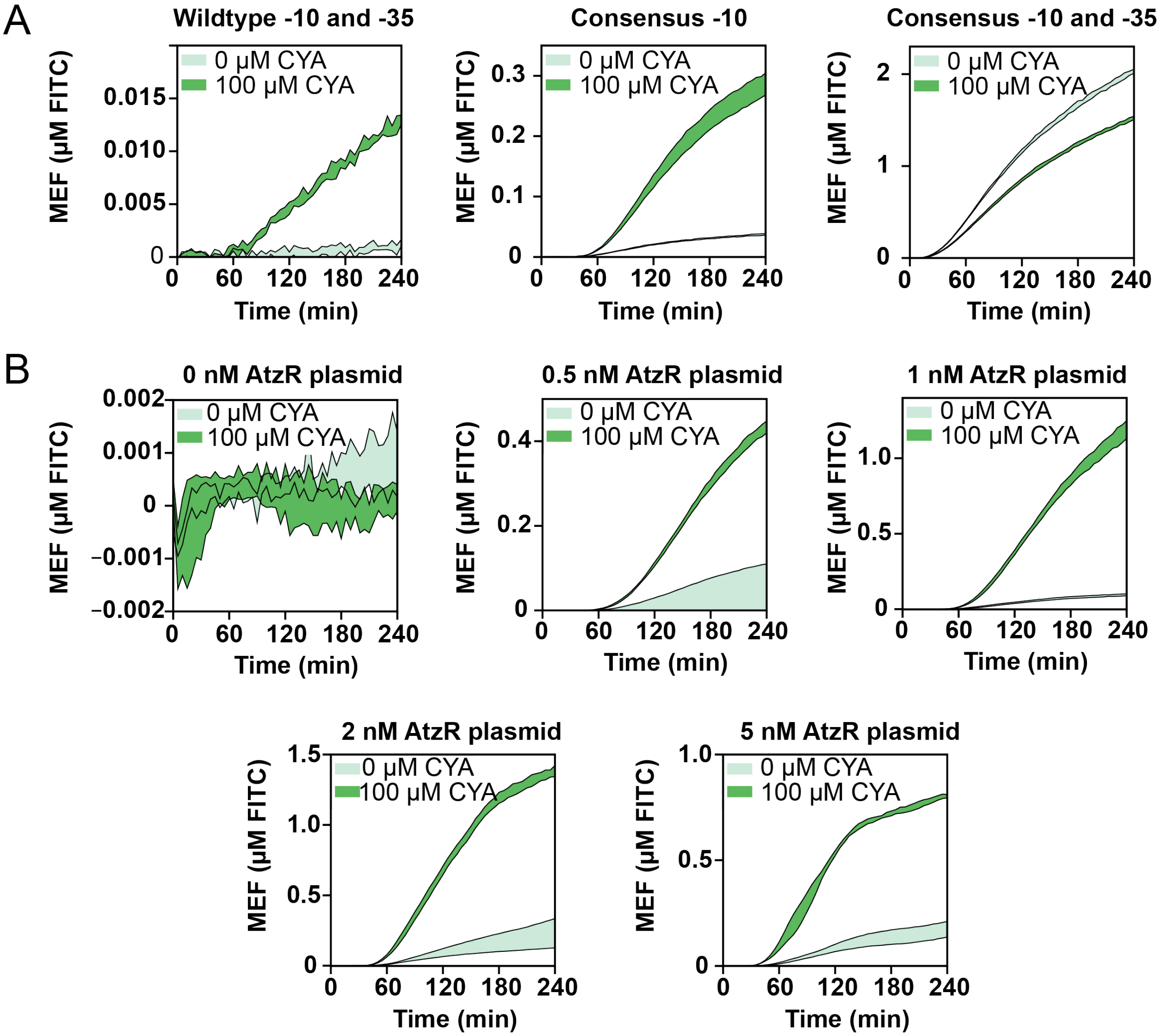


**Supplemental Figure 3**: **Kinetics of cyanuric acid reporter optimization in cell-free extracts**. (**A**) 4-hour time courses showing the differential kinetic responses of the data in **Fig. 3b** of the main manuscript for cell-free reporter designs. (**B**) 4-hour time courses showing the differential kinetic responses of the data in **Fig. 3c** of the main manuscript for AtzR plasmid concentration. Error shading represents one standard deviation from 3 technical replicates. Fluorescence is reported standardized to mean equivalent fluorescence (MEF) of fluorescein isothiocyanate based on a previously developed standard.

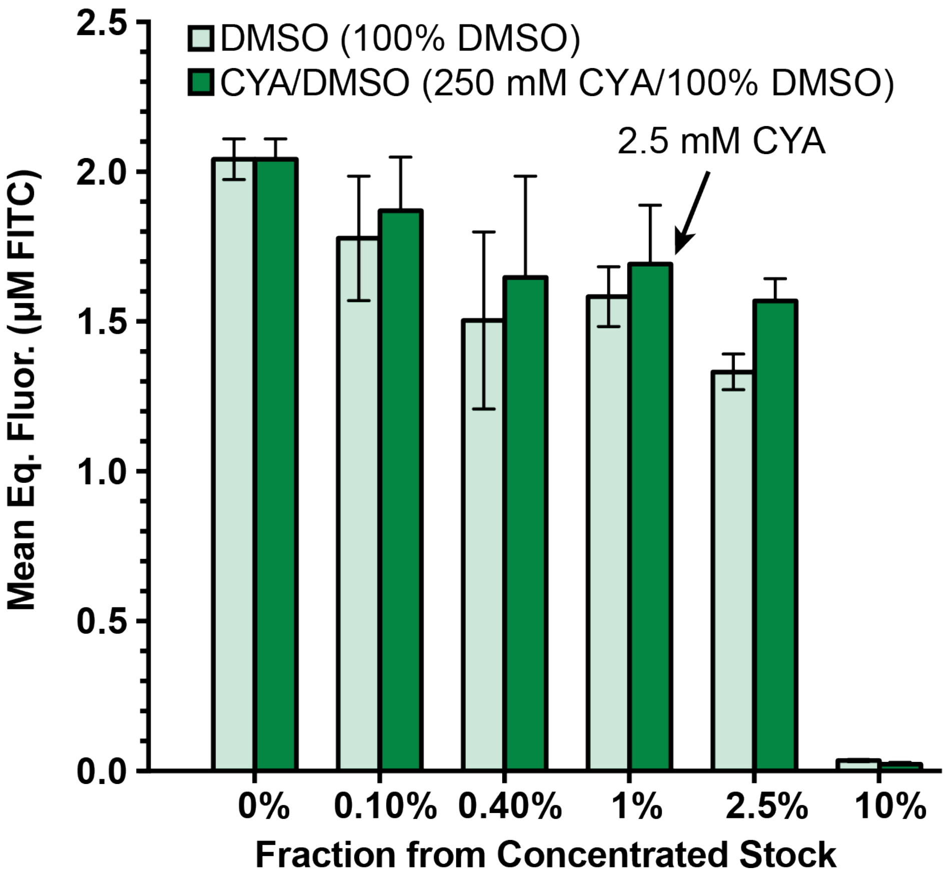


**Supplemental Figure 4: Cyanuric acid inhibition of cell-free protein synthesis**. A stock of either 250 mM cyanuric acid (dissolved in DMSO) or pure DMSO was serially diluted into a cell-free reaction constitutively expressing sfGFP without any transcription regulation. A value of 1% implies that the reaction was supplemented either with DMSO or 250 mM cyanuric acid (dissolved in DMSO), to a final volume fraction of 1%. Error bars represent 1 standard deviation from 3 technical replicates. Fluorescence is reported standardized to mean equivalent fluorescence (MEF) of fluorescein isothiocyanate based on a previously developed standard.

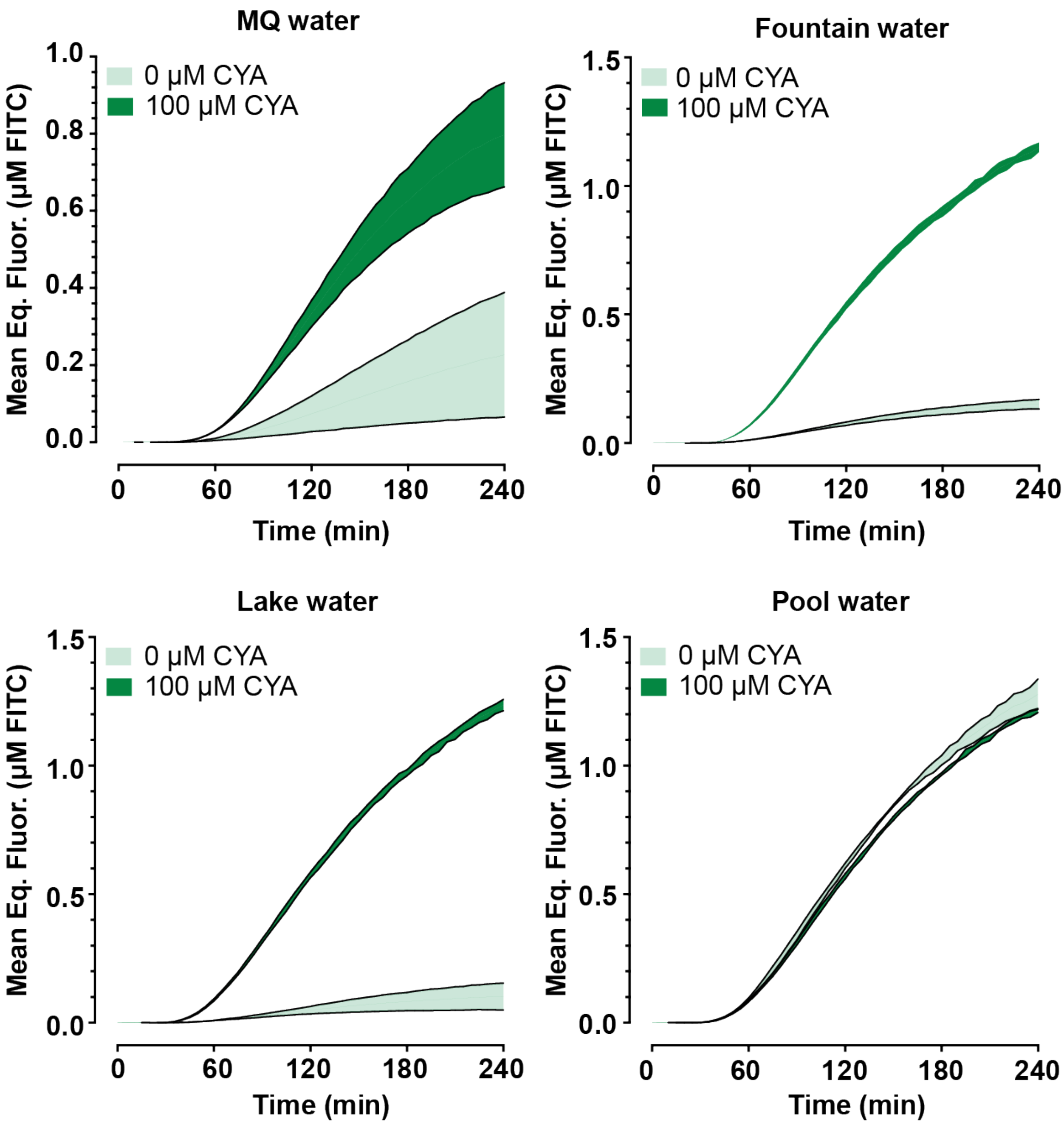


**Supplemental Figure 5: Kinetics of freeze-dried cell-free detection of cyanuric acid.** 4-hour time courses showing the differential kinetic responses of the data in **Fig. 4b** of the main manuscript for detecting MilliQ, fountain, lake, and pool water, with externally supplemented cyanuric acid. Error shading represent 1 standard deviation from 3 technical replicates. Fluorescence is reported standardized to mean equivalent fluorescence (MEF) of fluorescein isothiocyanate based on a previously developed standard.

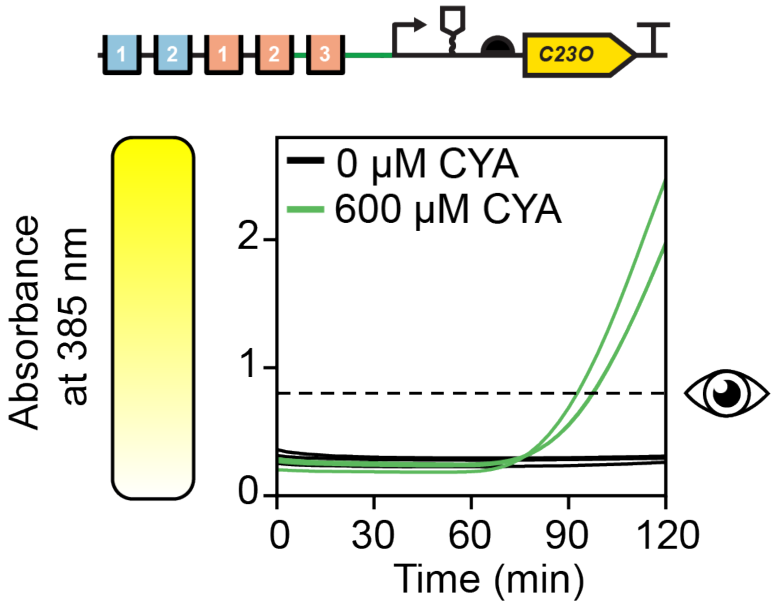


**Supplemental Figure 6: Kinetics of cell-free colorimetric reporter**. 2-hour time course showing the kinetic response, measured on a plate reader, of the C23DO reporter in response to high (600 µM) cyanuric acid or plain water. The eye icon to the right of the graph represents the estimated visible threshold of absorbance based on previous empirical measurements from our lab.[1]

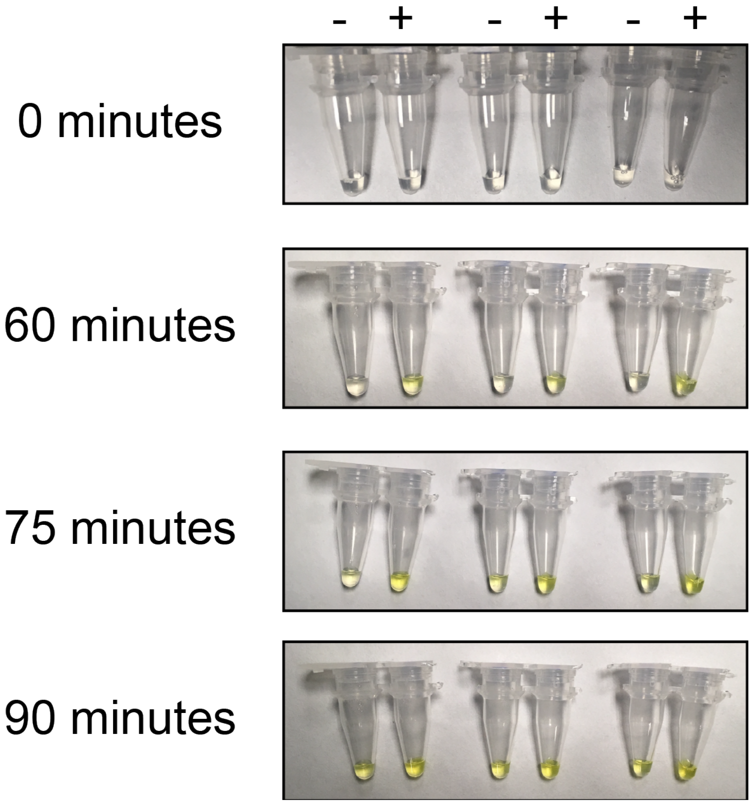


**Supplemental Figure 7: Kinetics for equipment-free visual detection of cyanuric acid.** Uncropped images from **Fig. 4c** of the main manuscript including technical replicates and an additional measured time point.

| Component | Cost/10 µL reaction | Cost/100 reactions |
|---|---:|---:|
| Postlysis processed extract | $0.010 | $0.97 |
| Phosphoenolpyruvate (PEP) | $0.017 | $1.74 |
| Nucleotide triphosphates (NTPs) | $0.007 | $0.75 |
| Midiprepped plasmid DNA | $0.006 | $0.57 |
| All other reagents | $0.008 | $0.81 |
| **Total** | **$0.048** | **$4.84** |

**Supplemental Table 1**: Estimated raw cost of a cell-free cyanuric acid sensor. Calculations were made based on 2019 commercial prices for commodity chemicals available from Sigma-Fisher and the referenced protocol[2]. The following additional assumptions were made: 3 mL of extract obtained per liter of culture; 100 µg plasmid DNA obtained per midiprep kit; 5 nM reporter plasmid and 1 nM transcription factor plasmid, and 2 mM catechol supplied to the reaction. Only raw materials costs (excepting labor and capital) were considered in this calculation.

| Plasmid name & architecture | Sequence 5’ to 3’ |
|---|---|
| pXL-P-intragenic (Intragenic-sfGFP-ColE1-KanR) | ATGCGAGTCAAAGCAAGATCGGTGCCGGATCGGCAC<br>CAGTTAGGTCGGAAAAAGGCGGCAGTCAAGTGCGCA<br>GGGCGGCGTTAAGCTTGAACGAAATGTTCTGCCTGG<br>GCGCAGTTGCGCCAGGCCGTGTAGTGACGTCGCTCG<br>GTGCATGTACAGGGAACAGCCATCCGTCCTATTAACC<br>TTTTTGAGAATTGCCAGAT |
| pXL-P-WT (WT-Bujard RBS-sfGFP-ColE1-KanR) | ATGCGAGTCAAAGCAAGATCGGTGCCGGATCGGCAC<br>CAGTTAGGTCGGAAAAAGGCGGCAGTCAAGTGCGCA<br>GGGCGGCGTTAAGCTTGAACGAA<br>GAATTCATTAAAGAGGAGAAAGGT |
| pXL-P1 (P1-Bujard RBS-sfGFP-ColE1-KanR) | ATGCGAGTCAAAGCAAGATCGGTGCCGGATCGGCAC<br>CAGTTAGGTCGGAAAAAGGCGGCAGTCAAGTGCGCA<br>GGGCGGCGTTATAATTGAACGAA<br>GAATTCATTAAAGAGGAGAAAGGT |
| pXL-P2 (P2-Bujard RBS-sfGFP-ColE1-KanR) | TCGGTGCCGGATCGGCACCAGTTAGGTCGGAAAAAG<br>GCGGCAGTCAAGTGCGCTTTTTGTACCTATAATAGATT<br>CATGATGA GAATTCATTAAAGAGGAGAAAGGT |
| pXL-P3 (P3-Bujard RBS-sfGFP-ColE1-KanR) | TCGGTGCCGGATCGGCACCAGTTAGGTCGGAAAAAG<br>GCGGCAGTCAAGTGCGCTTTTTGTACCTATATAGATTC<br>ATGATGA GAATTCATTAAAGAGGAGAAAGGT |
| pXL-P4 (P4-Bujard RBS-sfGFP-ColE1-KanR) | TCGGTGCCGGATCGGCACCAGTTAGGTCGGAAAAAG<br>GCGGCAGTCAAGTGCGCTTTTTGTACCTATAAAGATT<br>CATGATGA GAATTCATTAAAGAGGAGAAAGGT |
| pXL-P5 (P5-Bujard RBS-sfGFP-ColE1-KanR) | TCGGTGCCGGATCGGCACCAGTTAGGTCGGAAAAAG<br>GCGGCAGACAAGTGCGCTTTTTGTACCTATAATAGATT<br>CATGATGA GAATTCATTAAAGAGGAGAAAGGT |
| pXL-P6 (P6-Bujard RBS-sfGFP-p15a-KanR) | TCGGTGCCGGATCGGCACCAGTTAGGTCGGAAAAAG<br>GCGGCAGACAAGTGCGCTTTTTGTACCTATAATAGATT<br>CATGATGA GAATTCATTAAAGAGGAGAAAGGT |
| pXL-P7 (P7-Bujard RBS-sfGFP-ColE1-KanR) | TCGGTGCCGGATCGGCACCAGTTAGGTCGGAAAAAG<br>GCGGCAGACAAGTGCGCTTTTTGTACCTATATAGATT<br>CATGATGA GAATTCATTAAAGAGGAGAAAGGT |
| pXL-P8 (P8-Bujard RBS-sfGFP-ColE1-KanR) | TCGGTGCCGGATCGGCACCAGTTAGGTCGGAAAAAG<br>GCGGCAGACAAGTGCGCTTTTTGTACCTATAAAGATT<br>CATGATGA GAATTCATTAAAGAGGAGAAAGGT |
| pXL-P9 (P9-Bujard RBS-sfGFP-ColE1-KanR) | TCGGTGCCGGATCGGCACCTGACAGGTCGGAAAAAG<br>GCGGCTATAAGTGCGCATGATGA<br>GAATTCATTAAAGAGGAGAAAGGT |
| PXL-AtzR (apFab61-Bba_J61132-AtzR-SC101 (high copy)-SpecR) | TTGACAATTAATCATCCGGCTCGTTTAATAGATTCATT<br>AGAGTCTAGAGAAAGACAGGATTAACATGCAACACCT<br>GCGTTTCCTGCACTACATCGACGCGGTTGCGCGTTGC<br>GGTAGCATCCGTGCGGCGGCGGAGCAACTGCATGTT<br>GCGGCGAGCGCGGTGAACCGTCGTGTTCAAGATCTG<br>GAGTACGAACTGGGTACCCCGATCTTTGAGCGTCTGC |

| | |
|---|---|
| | CGCGTGGTGTGCGTCTGACCGCGGCGGGTGAACTGT<br>TTGTTGCGTATGCGCGTCGTCGTAACGCGGACCTGGA<br>ACAGGTGCAAAGCCAGATTCAAGATCTGAGCGGTATG<br>AAGCGTGGCCGTGTTACCCTGGCGGCGAGCCAGGCG<br>CTGGCGCCGGAGTTCCTGCCGCGTGTGATCCACGCG<br>TTTCAGGCGCAACGTCCGGGTATTGCGTTCGACGTGA<br>AAGTTCTGGATCGTGAACGTGCGGTGCTGGCGGTTAC<br>CGACTTTGCGGCGGATCTGGCGCTGGTGTTTAACCC<br>GCCGGACCTGCGTGGCCTGACCGTTATTGCGCAGGC<br>GCGTCAACGTATTTGCGCGGTGGTTGCGAGCGATCA<br>CCCGCTGGCGAAGCGTACCAGCCTGCGTCTGAAAGA<br>CTGCCTGGATTACCCGCTGGCGCTGCCGGACAGCAG<br>CCTGAGCGGTCGTAACGTGCTGGACGAGCTGTTTGAT<br>AAAAGCAGCGCGCGTCCGCGTCCGCAGCTGGTTAGC<br>AACAGCTATGAGATGATGCGTGGCTTCGCGCGTGAAA<br>CCGGTGGCGTGAGCTTTCAAATCGAAATTGGTGCGG<br>GCAGCACCGAGGGTGAAGTTGCGATCCCGATTGATG<br>AGCGTAGCCTGGCGAGCGGCCGTGTGGTTCTGGTTG<br>CGCTGCGTGAACGTGTGCTGCCGGTTGCGAGCGCGG<br>CGTTTGCGGAGTTCGTTGCGGGCAAGCTGGCGGATA<br>CCACCCATAATGCGACCTAA |
| pADS108 (promoter-RNA stability hairpin-RBS-sfGFP-TrrnB-ColE1-AmpR) | CATGCGAGTCAAAGCAAGATCGGTGCCGGATCGGCA<br>CCAGTTAGGTCGGAAAAAGGCGGCAGTCAAGTGCGC<br>AGGGCGGCGTTATAATTGAACGAAACGTCGACTCTCG<br>AGTGAGATTGTTGACGGTACCGTATTTTggatctaggagga<br>aggatct |
| pADS109 (promoter-RNA stability hairpin-RBS-sfGFP-TrrnB-ColE1-AmpR) | CATGCGAGTCAAAGCAAGATCGGTGCCGGATCGGCA<br>CCAGTTAGGTCGGAAAAAGGCGGTTGACAAGTGCGC<br>AGGGCGGCGTTATAATTGAACGAAACGTCGACTCTCG<br>AGTGAGATTGTTGACGGTACCGTATTTTggatctaggagga<br>aggatct |
| pADS114 (T7 promoter-RBS-AtzR-ColE1-KanR) | taatacgactcactatagggagaccacaacggtttccctctagaaataattttgttt<br>aactttaagaaggagatatacatATGCGCGCTCAACGTCCTAGT<br>GCGATTCAGGCAATTAGCATTCCTTCCTTTCTGTTACG<br>TCCTCCGATGCAACACCTGCGCTTTTTGCACTACATC<br>GACGCTGTCGCTCGCTGCGGATCCATCCGTGCTGCC<br>GCCGAGCAGCTTCATGTTGCAGCATCCGCGGTCAAC<br>CGTCGCGTTCAAGATTTAGAATATGAACTGGGTACCC<br>CAATCTTCGAACGTTTACCCCGCGGCGTTCGCCTGAC<br>CGCCGCGGGTGAACTGTTCGTCGCGTATGCGCGTCG<br>TCGTAATGCCGATTTAGAGCAGGTGCAATCGCAAATC<br>CAAGATTTGAGCGGCATGAAACGCGGCCGTGTCACG<br>CTGGCCGCCTCACAAGCTTTAGCCCCGGAATTCTTAC<br>CTCGCGTAATTCACGCGTTTCAGGCACAACGCCCGG<br>GCATCGCGTTCGACGTAAAGGTGCTGGATCGCGAGC<br>GTGCGGTTCTTGCGGTGACTGATTTTGCGGCAGATTT |

| | |
|---|---|
| | AGCTCTGGTGTTTAATCCCCCTGATTTACGCGGTCTTA<br>CGGTTATTGCACAAGCCCGCCAACGTATCTGTGCAGT<br>TGTGGCCTCGGACCACCCCCTTGCTAAACGTACTTCA<br>CTTCGCCTGAAGGACTGCTTGGACTACCCATTGGCCC<br>TTCCAGACTCGTCATTGAGCGGCCGCAATGTCTTAGA<br>TGAATTGTTTGATAAATCCTCAGCACGCCCACGTCCG<br>CAACTGGTCAGCAATTCATATGAGATGATGCGCGGCT<br>TCGCCCGTGAGACGGGGGGGTATCTTTCCAAATCG<br>AAATCGGGGCTGGTTCTACGGAGGGGGAAGTCGCCA<br>TTCCGATCGACGAACGTTCGCTGGCCTCGGGGCGTG<br>TAGTTCTGGTTGCACTGCGCGAACGCGTACTTCCTGT<br>CGCATCGGCCGCATTTGCCGAGTTCGTCGCAGGTAAA<br>TTAGCAGACACGACTCATAACGCAACCtaa |
| pADS115<br>(promoter-RNA stability hairpin-RBS-catecholase-TrrnB-ColE1-AmpR) | CATGCGAGTCAAAGCAAGATCGGTGCCGGATCGGCA<br>CCAGTTAGGTCGGAAAAAGGCGGCAGTCAAGTGCGC<br>AGGGCGGCGTTATAATTGAACGAAACGTCGACTCTCG<br>AGTGAGATTGTTGACGGTACCGTATTTTggatctaggagga<br>aggatctatgaacaaaggtgtaatgcgaccgggccatgtgcagctgcgtgtact<br>ggacatgagcaaggccctggaacactacgtcgagttgctgggcctgatcgag<br>atggaccgtgacgaccagggccgtgtctatctgaaggcttggaccgaagtgga<br>taagttttccctggtgctacgcgaggctgacgagccgggcatggattttatgggttt<br>caaggttgtggatgaggatgctctccggcaactggagcgggatctgatggcat<br>atggctgtgccgttgagcagctacccgcaggtgaactgaacagttgtggccgg<br>cgcgtgcgcttccaggccccctccgggcatcacttcgagttgtatgcagacaag<br>gaatatactggaaagtggggtttgaatgacgtcaatcccgaggcatggccgcg<br>cgatctgaaaggtatggcggctgtgcgtttcgaccacgccctcatgtatggcga<br>cgaattgccggcgacctatgacctgttcaccaaggtgctcggtttctatctggccg<br>aacaggtgctggacgaaaatggcacgcgcgtcgcccagtttctcagtctgtcg<br>accaaggcccacgacgtggccttcattcaccatccggaaaaaggccgcctcc<br>atcatgtgtccttccacctcgaaacctgggaagacttgcttcgcgccgccgacct<br>gatctccatgaccgacacatctatcgatatcggcccaacccgccacggcctca<br>ctcacggcaagaccatctacttcttcgacccgtccggtaaccgcaacgaagtgt<br>tctgcgggggagattacaactacccggaccacaaaccggtgacctggaccac<br>cgaccagctgggcaaggcgatcttttaccacgaccgcattctcaacgaacgatt<br>catgaccgtgctgacctga |
| pJBL6853_PatzDEF_sfGFP<br>(promoter-RNA stability hairpin-RBS-sfGFP-TrrnB-ColE1-AmpR) | CATGCGAGTCAAAGCAAGATCGGTGCCGGATCGGCA<br>CCAGTTAGGTCGGAAAAAGGCGGCAGtCAAGTGCGCA<br>GGGCGGCGTTAAGCTTGAACGAAACGTCGACTCTCG<br>AGTGAGATTGTTGACGGTACCGTATTTTggatctaggagga<br>aggatct |

**Supplemental Table 2**: **Plasmids used in this study.** Features are highlighted in colors according to their names for each plasmid.
